# No evidence that mate choice in humans is dependent on the MHC

**DOI:** 10.1101/339028

**Authors:** Mircea Cretu-Stancu, Wigard P. Kloosterman, Sara L. Pulit

**Author notes:** **Corresponding authors:** Dr. Sara L. Pulit, Department of Genetics, University Medical Center Utrecht, Heidelberglaan 100, 3584 CX Utrecht, Utrecht, The Netherlands,; Dr. Wigard Kloosterman, Department of Genetics, University Medical Center Utrecht, Heidelberglaan 100, 3584 CX Utrecht, Utrecht, The Netherlands.

## Abstract

A long-standing hypothesis in biology proposes that various species select mates with a major histocompatibility complex (MHC) composition divergent from their own, so as to improve immune response in offspring. However, human and animal studies investigating this mate selection hypothesis have returned inconsistent results. Here, we analyze 239 mate-pairs of Dutch ancestry, all with whole-genome sequence data collected by the Genome of the Netherlands project, to investigate whether mate selection in humans is MHC dependent. We find no evidence for MHC-mediated mate selection in this sample (with an average MHC genetic similarity in mate pairs (Qc) = 0.829; permutation-based p = 0.703). Limiting the analysis to only common variation or considering the extended MHC region does not change our findings (Qc = 0.671, p = 0.513; and Qc = 0.844, p = 0.696, respectively). We demonstrate that the MHC in mate-pairs is no more genetically dissimilar (on average) than a pair of two randomly selected individuals, and conclude that there is no evidence to suggest that mate choice is influenced by genetic variation in the MHC.

**Author summary:** Studies within various animal species have shown that the genetic content of the major histocompatibility complex (MHC) can influence mate choice. Such mate selection would be advantageous, as mating between individuals with different alleles across MHC genes would produce offspring with a more diverse MHC and therefore possess improved immune response to various pathogens. Studies of the influence on the MHC in human mate selection have been far less conclusive. Two studies of MHC-dependent mate selection performed on SNP data collected as part of the HapMap Consortium returned conflicting results: the first study reported significantly different MHC variation between mate pairs, and the second report refuted this claim. Here, we analyze a dataset comprised of 239 whole-genome sequenced Dutch mate pairs, a sample set an order of magnitude larger than the HapMap data and containing denser characterization of genetic variation. We find no evidence that the MHC influences mate selection in our population, and we show that this finding is robust to potential confounding factors and the types and frequencies of genetic variants analysed.

## Introduction

The extended major histocompatibility complex (MHC) spans an approximately 7-megabase region on chromosome 6 in humans. The region codes for a series of proteins critical to acquired immune function as well as olfactory genes [1]. Additionally, the MHC contains extensive genetic diversity [2,3], much more so than other regions of the genome; within the human population, the MHC contains thousands of different alleles and haplotypic combinations spanning the frequency spectrum. Genome-wide association studies (GWAS) have identified a plethora of genetic variants in the region associated to a host of diseases [4], both with and without previously-described roles for immune function [5–10].

Some biological studies have proposed that, beyond the direct role in immune function, the MHC may influence mate selection in vertebrate species. Increased MHC diversity is evolutionarily advantageous, as it improves immune response to a wider range of pathogens [11,12]. A number of studies in (non-human) animals indicate that some species of mice, birds, and fish, preferentially mate to maintain or increase MHC diversity [13–17]. For example, studies in sticklebacks [18] indicate that MHC-based mate selection helps to optimize copy number of particular MHC loci between mates. In mice, increased MHC dissimilarity between mates increases diversity of amino acid substitutions within binding-pockets of specific HLA molecules [19,20]. Many of these studies suggest that the observed MHC-dependent mate selection is mediated by the olfactory system, either through detectable residues that mates can smell [21], or because olfactory receptor genes are often found to cluster in close genomic proximity to the MHC [3].

Evidence for MHC-dependent mate selection in humans is far less conclusive. A study of 411 couples from the Hutterite population, a population isolate in North America, performed HLA typing across all couples and found that couples had more MHC diversity than expected under random mating [22]. Two additional studies, of 200 Amerindian couples [23] and 450 Japanese couples [24], respectively, concluded that the differences between the HLA-types of real couples were not significantly more different than the HLA types of random pairs of individuals. Finally, additional work has investigated whether the remnants of degraded HLA proteins end up in sweat, urine or saliva and can therefore be detected by potential mates through scent. To test the hypothesis that MHC-dependent mate selection in humans is mediated through olfactory processes, researchers have performed so-called ‘sweaty t-shirt’ experiments, and shown that females indicate an odor preference towards men that carry divergent HLA alleles relative to their own [25,26].

Studies of genetic variation (beyond the classical HLA types) in humans have sought to provide clarity as to whether humans do indeed select mates, at least in part, such that diversity across the MHC increases in offspring. An initial analysis of array-based SNP genotyping data (variation with minor allele frequency (MAF) > 5%) assembled by the HapMap 2 Consortium [27] examined 30 European-ancestry mate pairs and 30 African-ancestry mate pairs and reported evidence of dissimilar MHC variation in couples of European descent (p = 0.015) [17]. Conversely, no such effect was observed in the African-ancestry sample (p = 0.23) [17]. A subsequent analysis in the same Hapmap Phase 2 European-ancestry data, but including an additional 24 European-ancestry mate-pairs genotyped as part of HapMap Phase 3 [28], failed to replicate the initial finding [29]. This second analysis demonstrated that the low sample size of the initial analysis (making the study sensitive to small changes in parameter choices) and failure to correct for multiple testing explained the initial report. Neither analysis of the 24 new mate-pairs nor joint analysis of all 54 available European-ancestry mate pairs revealed increased MHC dissimilarity in mates (p = 0.351 and p = 0.143, respectively).

Here, we aim to test whether human mate pairs are indeed more dissimilar across the MHC, using a sample set that represents an order-of-magnitude increase over the initial reports. Specifically, we test the hypothesis that MHC variation is discordant between couples by analyzing a dataset of 239 unrelated Dutch mate pairs, whole-genome sequenced as part of the Genome of the Netherlands (GoNL) project [30]. The density and resolution of the whole-genome sequence data allow us to test for discordant MHC variation in mate pairs with respect to (a) common variation only (MAF > 1%); (b) the full frequency spectrum of genetic variants, including single nucleotide variants and short insertions and deletions; and (c) imputed amino acids and human leukocyte antigen (HLA) types within the MHC [31].

## Results

### Reproducing the initial HapMap analysis

We first sought to reproduce the finding of MHC-dependent mate selection in humans reported from an analysis of common variation in the Hapmap Phase 2 data [17], with the goal of not only replicating results but also aligning methodologies. The previous analysis used 30 trios of Northern- and Western-European ancestry living in Utah, USA (called the CEU sample) and 30 trios collected from the Yoruba population in Ibadan, Nigeria (called the YRI sample) [27,32,33] to evaluate MHC genetic dissimilarity in mate pairs. After reproducing the quality control procedures from the initial analysis as closely as possible (**Materials and Methods**), 27 CEU and 27 YRI mate-pairs remained for analysis (**Table 1**).

**Table 1.**
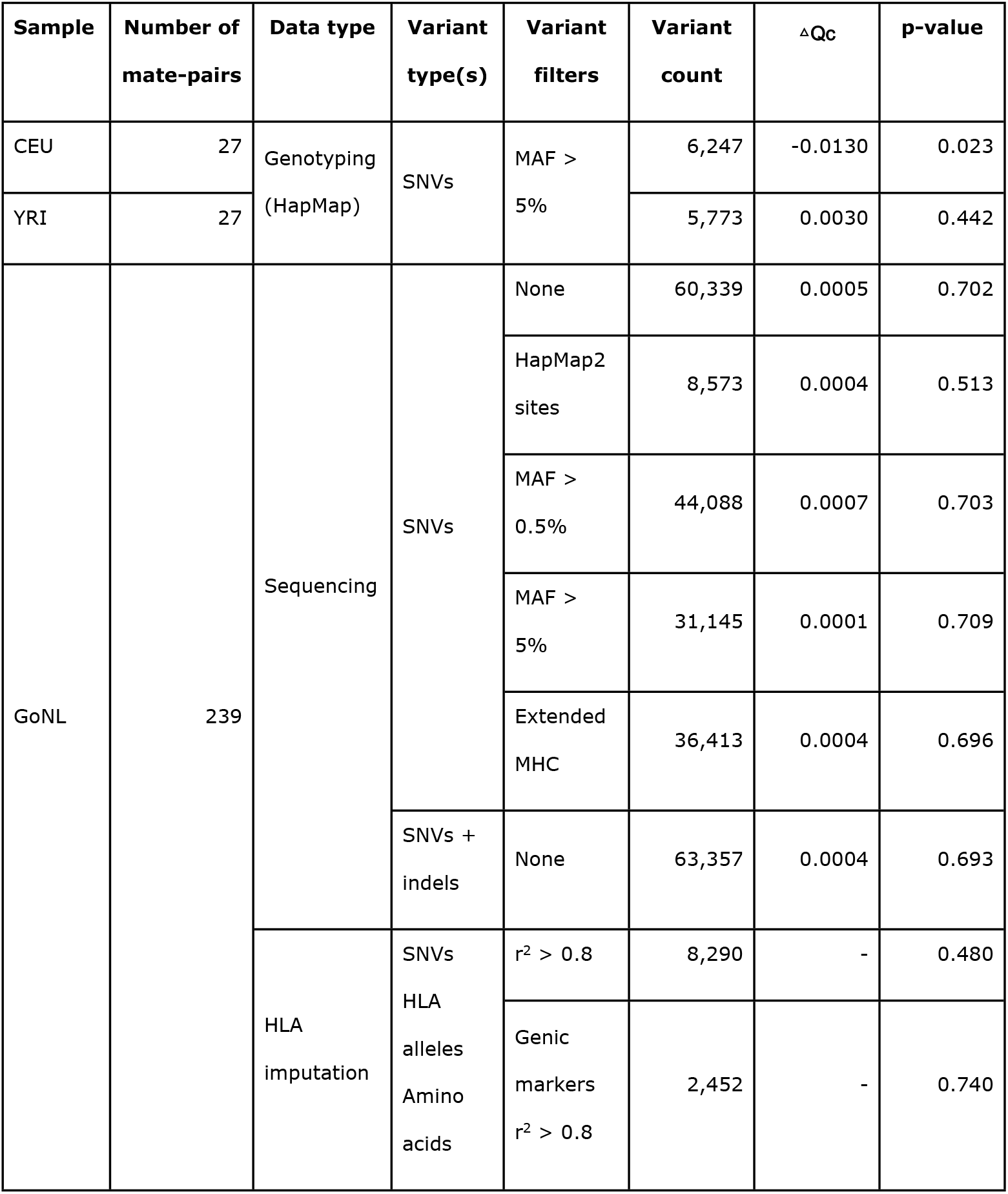
Samples and variants used in analysis. To investigate whether mate selection is MHC-dependent, we analyzed three sample groups: Utah residents with Northern and Western European ancestry (CEU); Yorubans from Ibadan, Nigeria (YRI); and mate-pairs in the Genome of the Netherlands (GoNL) project. The number of mate pairs indicates the number of pairs available after sample quality control. We performed our analysis in common polymorphisms (minor allele frequency (MAF) > 0.05) or common- and low-frequency single nucleotide variants (SNVs, with MAF > 0.5%), as well as including indel variation, where available. For imputed data, we kept only well-imputed data, based on the Beagle imputation quality metric (r^2^ > 0.8). We additionally restricted the set of variants to only variants within the classical HLA genes including amino acid substitutions, single nucleotide polymorphisms (SNP), insertions and/or deletions (indels) and classical HLA-types (‘genic markers’).

We used the same measure for genetic similarity between two individuals as defined in the initial report: Qc, defined as ‘the proportion of identical genotypes (at variant positions)’ [17] between mate pairs (**Materials and Methods**). We compared the average similarity across real couples to the average similarity across randomly generated mate pairs (created by randomly drawing a male and a female from the sample) and obtained results that are close, but not identical to, the initial report (**Figure 1**). We calculated the difference between average genetic similarity across all true mate pairs and average genetic similarity across permuted mate pairs (i.e., average Qc across a null distribution; **Figure 1**) to explicitly quantify how genetic similarity in true mate pairs deviates from the null distribution. We call this metric ΔQc. We found that the CEU mate pairs demonstrated nominally-significant (p < 0.05) genetic dissimilarity across the MHC compared to permuted mate pairs (ΔQc = -0.013, 2-sided p = 0.023), while mate-pairs in the YRI samples indicated no such relationship (ΔQc = 0.003, 2-sided p = 0.442). Genome-wide, CEU mate pairs showed no pattern of genetic similarity or dissimilarity (ΔQc = -0.008, 2-sided p = 0.100) while YRI mate-pairs showed a pattern of genome-wide similarity (average Qc = 0.011, 2-sided p < 10^-6^), consistent with the original report [17].

**Figure 1.**
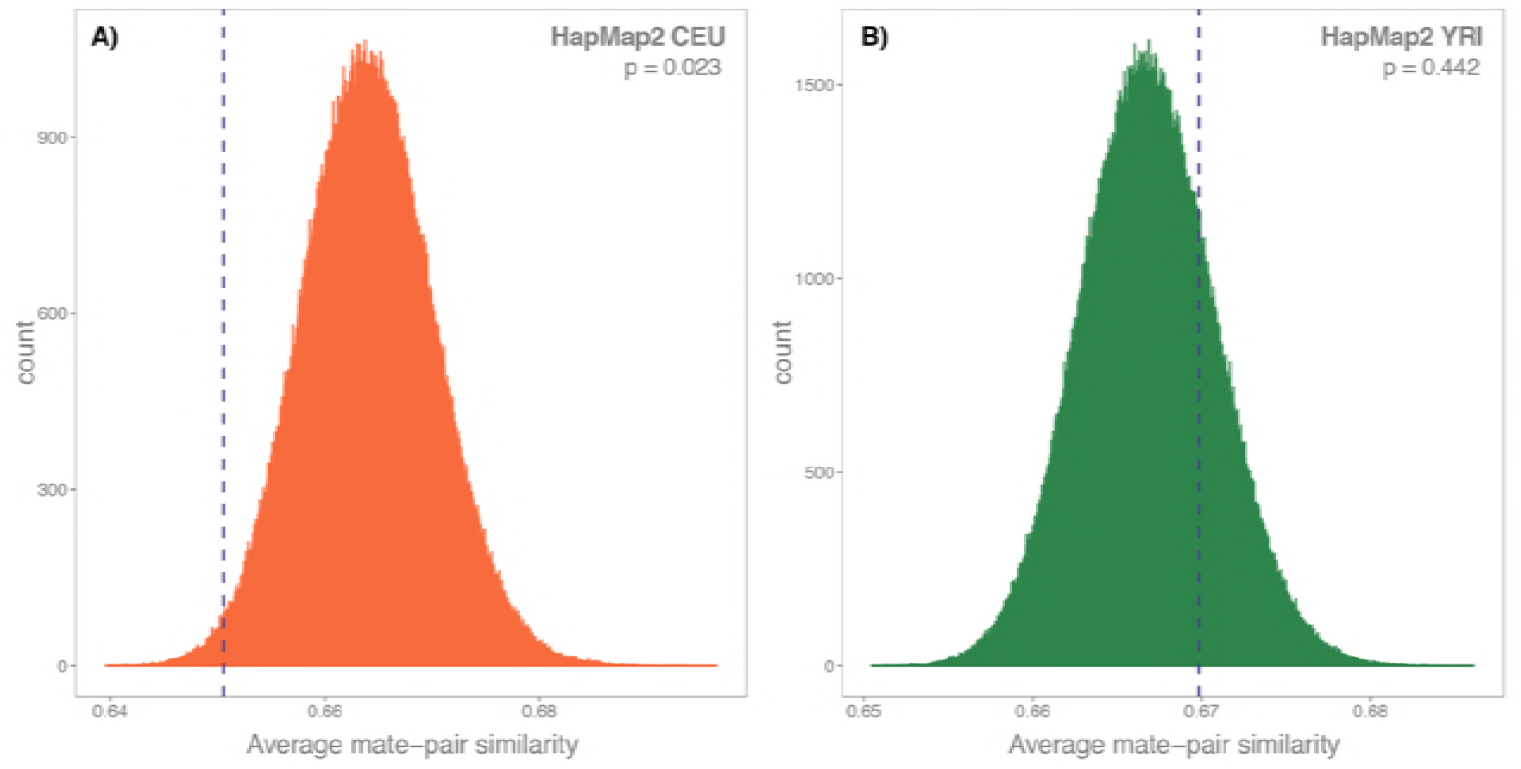
Genetic similarity across mate pairs in the HapMap 2 data. The distributions represent the null distribution of average MHC similarity (Qc), across randomly permuted mate pairs from each of the HapMap 2 populations tested (CEU: European samples of Northern and Western descent, orange; YRI: Yorubans in Ibadan Nigeria, green). The average MHC similarity in true mate pairs is marked by the blue dotted line. All p-values are based on 1,000,000 permutations and delta Qc (ΔQc) is the difference between the average real-couple similarity and the average of the distribution or random mate-pair permutations. **(A)** Permutation of the 27 QC-passing HapMap 2 CEU couples. ΔQc = -0.013, 2-sided p = 0.023. **(B)** Permutations of the 27 QC-passing HapMap 2 YRI couples. ΔQc = 0.003, 2-sided p = 0.442.

### Testing MHC-specific genetic dissimilarity in the Genome of the Netherlands

Next, we sought to test if there was evidence for MHC-dependent mate selection in mate pairs collected as part of the Genome of the Netherlands (GoNL) project [30]. GoNL is comprised of Dutch-ancestry trios (confirmed by principal component analysis [30]) drawn from 11 of the 12 provinces of the Netherlands and whole-genome sequenced at ~14x average coverage on the Illumina HiSeq 2000 [30]. After data quality control and processing in the original project [30], the GoNL dataset contained 248 mate pairs. Because relatedness is a primary confounder for genetic similarity estimations, we calculated sample relatedness in Plink [34] and removed an additional 9 mate pairs with pi-hat > 0.03125 (a threshold corresponding to 5th-degree relatedness; **Materials and Methods**). After this additional quality control, 239 mate pairs remained for analysis. We analyzed the GoNL data (http://www.nlgenome.nl/, see Code and Data Release in **Materials and Methods**) from Release 5 of the project, which includes single-nucleotide variants (SNVs) and short (< 20bp) insertions and deletions (indels; **Table 1**).

To test for MHC-dependent mate selection in GoNL, we extracted the MHC (chromosome 6, 28.7 - 33.3Mb on build hg19), calculated Qc across all true GoNL mate pairs, and performed the same permutation scheme as in the HapMap analysis, randomizing the mate pairs, recalculating the average Qc across these randomly-constructed pairs, and finally calculating ΔQc. All p-values are 1-sided, testing the hypothesis of genetic dissimilarity, unless otherwise stated. Our results showed no evidence for MHC-dependent mate selection (ΔQc = 0.0005, permutation p = 0.702, **Figure 2**). Restricting our analyses to common- and low-frequency SNPs (MAF > 0.5%) or common SNPs only (MAF > 5%) did not change our results (**Table 1**, **Supplementary Figures 1** and **2**), nor did restricting the analysis specifically to the ~2M common SNPs genotyped in HapMap 2 or including the set of ~2M indels sequenced in GoNL into the analysis (**Table 1** and **Supplementary Figure 3**). To test the hypothesis that MHC mating is mediated through olfactory sensory pathways, as hypothesized previously [25,26], we performed the same analysis using an extended definition of the MHC (26.6Mb - 33.3Mb on hg19), which includes a dense cluster of 36 olfactory receptor genes upstream of the HLA Class I region [3]. We observed no statistically significant effect (**Table 1**, and **Supplementary Figures 4** and **5**).

**Figure 2.**
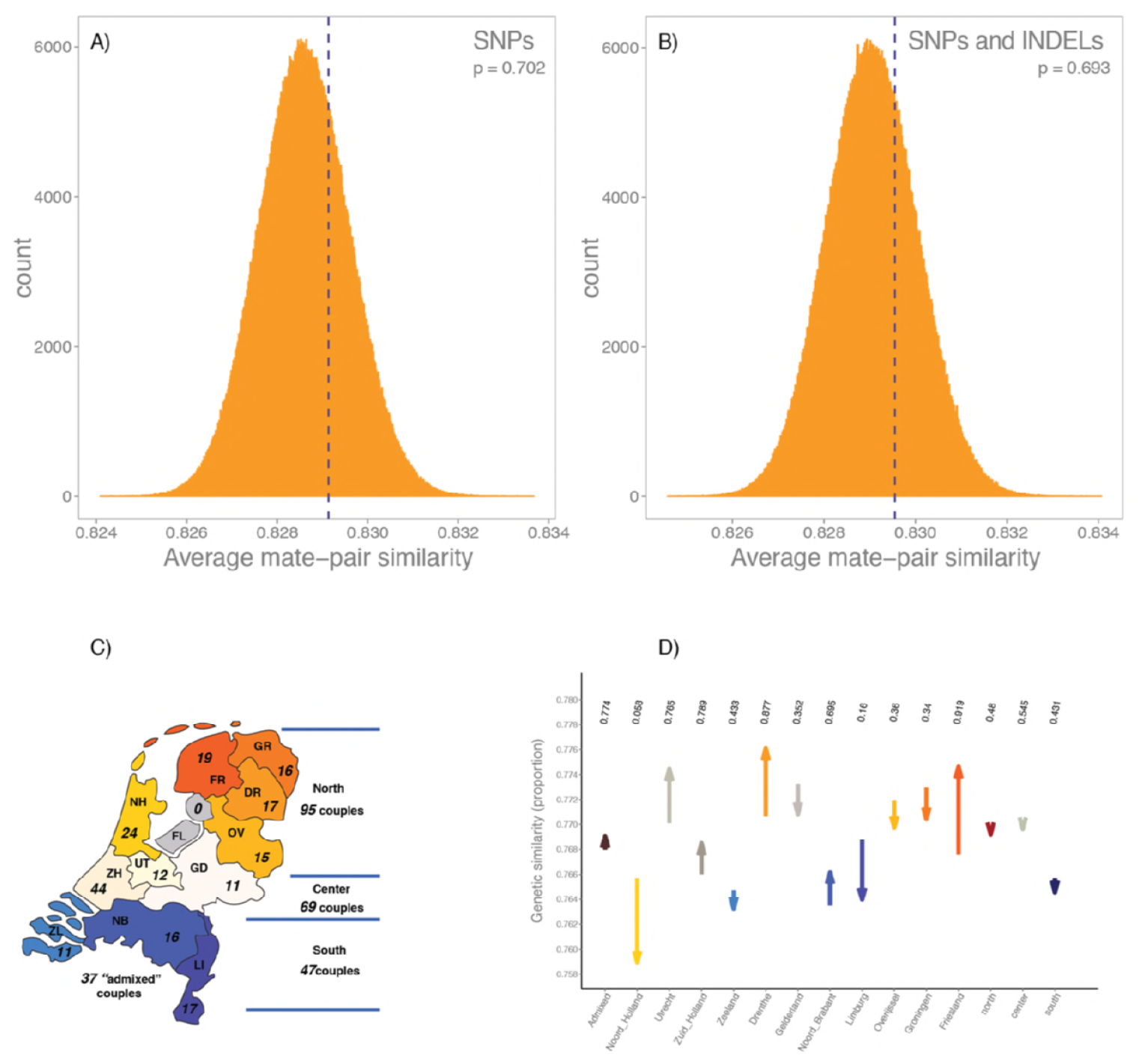
Genetic similarity across the MHC for 239 Dutch-ancestry mate-pairs. Panels **(A)** and **(B)** show the null distribution (histograms) of average mate-pair genetic similarity of permuted (i.e., non-real) male-female pairs. We performed a total of 1,000,000 permutations to generate the distribution. The average genetic similarity across 239 real mate pairs is represented with a blue vertical dotted line. **(A)** Genetic similarity measured across all biallelic variants within the MHC (p = 0.702). **(B)** Genetic similarity measured across all biallelic variants and insertions/deletions (indels) in the MHC (p = 0.693). **(C)** The GoNL samples were drawn from 11 of the 12 Dutch provinces. Here, we indicate the number of true mate-pairs available for analysis where both members of the mate-pair come from the same geographic region. These three geographic regions (north, center, and south) are derived from previously-performed population genetic analyses of the GoNL data. **(D)** Genetic similarity of mate-pairs, split by province. The arrows start at the average genetic similarity of permuted (i.e., null) mate pairs and stops at the average genetic similarity across true mate-pairs. Corresponding, one sided p-values for the genetic dissimilarity within couples are marked above.

Though the Netherlands is geographically small and densely populated, both common and rare variation in the GoNL data indicate geographic clustering [30,35–37]. We therefore investigated whether population stratification may explain the discordance between our results and the previous report of MHC-dependent mate selection in humans [17]. We performed genetic similarity analyses in the samples split into three geographic regions (“north,” “middle,” and “south” as determined by an identity-by-descent analysis [30]), as well as by province. Subsetting by region or province revealed no evidence for subpopulation-specific MHC-dependent mate selection (**Figure 2**). Additionally, accounting for sample ancestry using principal components (**Materials and Methods**) left our results unchanged (p = 0.78).

Lastly, we used SNP2HLA [31] to impute 2- and 4-digit HLA alleles, amino acids and SNPs (**Materials and Methods**) into the GoNL samples as a means of evaluating genetic (dis)similarity across imputed HLA types. Given that the dosages output from SNP2HLA are phased, we used the Pearson’s correlation (r) across the imputed allele dosages to calculate genetic similarity (instead of the Qc metric). We found no evidence for MHC-dependent mate selection either across all imputed markers (p = 0.48, **Table 1**) or by restricting the correlation calculation to only those variants, amino acids, and HLA types within the classical HLA Class I and II gene bodies (and thus more likely to have functional effect; p = 0.74, **Table 1**).

Until this point, we had established a null distribution by permuting mate pairs and calculating genetic similarity. To generate an alternative null model for comparison, we randomly sampled 10,000 regions from the genome that either matched the MHC by size (i.e., total span of the region) or by number of variants contained within the region (regardless of the total linear span of the region capturing those markers). For each permutation, we randomly selected the region, computed Qc (averaged across the 239 true mate-pairs) and counted the number of times the mean Qc was as or more dissimilar than that observed in the MHC. We observed no statistically significant difference, after accounting for multiple testing, when selecting regions based on genomic size or total number of markers in the region, after accounting for multiple testing (one-sided p = 0.08 and 0.02, respectively).

## Discussion

Using the whole-genome sequencing data of 239 mate pairs, we have performed, to our knowledge, the most comprehensive investigation of MHC-dependent human mate selection to date. The Genome of the Netherlands resource provided both an increased sample size compared to previous efforts [17,29] and high density genetic variation data, allowing for analyses of rare variants, indels, and imputed HLA types. However, despite the size and genomic resolution of the data, our results indicate no evidence for MHC-dependent mate selection in humans. We performed further analyses to investigate the potential effects of geographical clustering of rare variants [30,35], but the results left our results and interpretation unchanged.

Notably, our results are inconsistent with an initial investigation of MHC-dependent mate selection using genome-wide genetic variation data [17]. Though these previous findings do not align with our own, the initial report of MHC-dependent mate selection in humans was likely too small (N = 60) to draw conclusive results. Further, potential confounders, including cryptic relatedness and inbreeding amongst the studied samples, along with a lack of multiple testing correction, all likely contributed to this initial positive finding, subsequently contradicted in follow-up analyses of the same samples [29]. By interrogating a larger sample size, more stringently removing samples for relatedness and inbreeding, and performing analyses that account for potential population stratification, we believe our results provide more robust information as to whether mate selection in humans is influenced, at least in part, by individuals’ genetic composition across the MHC. Additionally, our results are consistent with investigation of MHC-dependent mate selection using HLA types in similarly-sized sample sets [23,24].

While our results indicate that human mate selection is independent of genetic variation in the MHC, a number of studies examining genetic variation and complex traits have found a plethora of positive evidence for assortative mating in humans based on non-MHC genetic factors. Previous studies have shown that human mate choice is associated to quantitative features (such as height) [38], to socioeconomic factors and risk for multifactorial disease [39–41]. A recent analysis in > 24,000 mate pairs, drawn from a number of cohorts including the UK Biobank [42] and 23andMe, focused on genomic loci associated to a number of multifactorial traits and found significant correlation between spouses at loci associated to height and body mass index [43]. By building a genetic predictor in one member of a spousal couple and applying it in the second member, the study also revealed varying degrees of spousal correlation at loci associated to waist-to-hip ratio, educational attainment, and blood pressure [43] in 7,780 couples from the UK Biobank. These correlations represent only a small slice of the numerous factors — both genetic and environmental — that contribute to mate selection in the human population. Importantly, however, these observations are correlative; the extent to which these associations are potentially causative remains to be explored.

Though our analysis offers several improvements over previous analyses examining MHC-dependent mate selection, several limitations remain. First, as highlighted by the assortative mating studies discussed above, our sample size may not be large enough to detect a more modest signal for MHC-dependent mate selection, if such a phenomenon exists. Mate selection is likely influenced by a host of hundreds, if not thousands, of factors, all of which likely have modest effect. Therefore, analysis of 239 samples may not be sufficiently well powered to detect such an effect. Further, while we have used permutations of mate pairs to establish a null distribution to which we can compare true mate-pair genetic similarity, this distribution may not be sufficiently informative to detect MHC-dependent effects. Indeed, the authors of the initial analyses [17] reported similar difficulties establishing a null comparator: they sought to additionally use genome-wide genetic similarity as a basis of comparison for MHC similarity, but observed higher genome-wide similarity in YRI samples compared to the CEU [17]. Given the uniqueness of the MHC, from its gene density and extensive linkage disequilibrium to its high genetic diversity, finding a genomic region with similar properties to use as a null comparator is essentially impossible; permutations of real mate pairs into random pairs, while not ideal, is likely the best null distribution for this experiment. Additionally, our analysis only examines one ancestral population. Analyses extended into other (non-European) samples may result in different findings.

Untested here is the hypothesis that preferential mating may favour specific combinations of HLA alleles that collectively result in an ‘optimal’ number of antigens that can be presented to T cell receptors. Previous studies indicate that this phenomenon may occur, specifically across Class I classical HLA genes [44], and may provide an alternative mechanism for MHC-mediated mate selection. Given the number of HLA allele combinations that would need to be constructed and analyzed to test such a hypothesis, power (after multiple test correction) would be vanishingly small. We therefore have not tested this specific hypothesis. However, additional information regarding gene function may make testing this hypothesis feasible in the future.

Despite these limitations, our analysis represents an improved investigation of MHC-dependent mate selection, through interrogated sample size as well as in the spectrum of genetic variation tested. Our data indicate no MHC-mediated preferential mating patterns in our European-ancestry sample. While MHC-mediated preferential mating has been reported in non-human animal models, such a mechanism in humans is either absent or may be one of many subtle contributors to mating patterns and behaviours.

## Materials and methods

### Code and data release

Individual-level data generated by the Genome of the Netherlands Project can be accessed through an application, available here: http://www.nlgenome.nl/. We provide code for this project at the following GitHub repository: https://github.com/mcretu-umcu/matingPermutations.

### Ethics Statement

All participants provided written informed consent as part of the Genome of the Netherlands project (http://www.nlgenome.nl/), and each biobank was approved by their respective institutional review board (IRB).

### Quality control of HapMap and Genome of the Netherlands data

Related samples, by definition, are more likely to share more genetic variation compared to two unrelated individuals. To ensure that relatedness was not confounding our analyses, we performed basic quality control (QC) in the CEU, YRI and Genome of the Netherlands (GoNL) sample sets separately. The initial HapMap 2 analysis [17] filtered related couples by looking at the normalized Qc measure and defining outliers. We used the identity-by-descent (IBD) estimates, computed with Plink 1.9 [45] using the --genome command. Though this approach differs from the initial analysis, using IBD estimates are an established means for identifying related samples using genetic variation data.

To estimate relatedness, we first used Plink 1.9 to assemble a set of high-quality SNPs with minor allele frequency (MAF) > 10% and genotyping missingness < 0.1%. We pruned this set of SNPs at a linkage disequilibrium (r^2^) threshold of 0.2. Additionally, we removed SNPs in the MHC, lactase (*LCT*) locus on chromosome 2, and in the inversions on chromosomes 8 and 17 (genomic coordinates in **Supplementary Table 1**). We calculated relatedness (--genome in Plink) across all individuals in the CEU and YRI mate pairs. We discarded three mate pairs (N = 6 samples) from the CEU sample and three mate pairs (N = 6 samples) from the YRI sample. We defined relatedness as pi-hat > 0.05 (i.e., shared 1/20th of the genome), close to the 1/22nd threshold used by Derti *et al*. [29]. Our filtering produced nearly identical results to the initial analyses (Supplementary Text S2 of [29]). Due to our slightly more stringent cutoff threshold, we additionally exclude the related pair of samples NA12892 and NA06994.

We filtered for relatedness in GoNL in an identical manner. We used a more stringent cryptic relatedness threshold of pi-hat > 0.03125, corresponding to 5th-degree relatives. We discarded 9 couples from our analysis, leaving 239 QC-passing mate pairs.

### Calculating genetic similarity in mate pairs

We define genetic similarity across a mate pair (called Qc, per the initial report [17]) as the proportion of variants that are identical across a pair of individuals. Homozygous genotypes comprised of the same alleles (e.g., AA in sample 1 and AA in sample 2) are considered 100% similar; heterozygous genotypes (e.g., AB in both samples) are considered 50% similar, as they could have either the same or opposite phase; and all other combinations are considered 0% similar.

We note that in the initial report [17], genetic similarity was defined as: R = (Qc - Qm)/(1-Qm), where Qm is the average genetic similarity across all possible mate-pairs (real and permuted) that can be constructed in the sample. We note that the R measure is a linear transformation of Qc measure, as Qm is a constant for the analyzed sample. Further, Qm is not an unbiased estimate of the average genetic similarity within random mate-pairs for two reasons: (1) because it includes both real mate-pairs and female-male pairs constructed by selecting two random individuals in the dataset; and (2) because the sample pairs over which Qm is averaged are not independent (i.e., the same individual is paired with all possible matches and thus considered multiple times when computing Qm). We therefore perform all our analyses using only the Qc measure of genetic similarity.

### Replicating the original HapMap analysis

The HapMap 2 genotyping data is publicly available [27,32,33] and includes a total of 3,965,296 single nucleotide polymorphisms (SNPs). We extracted the MHC region (29.7 - 33.3Mb on chromosome 6, build hg18, as defined in the original analysis) from each population separately: people of Northern and Western European ancestry (the CEU) and Yorubans from Ibadan, Nigeria (YRI). We performed these analyses in 27 CEU mate-pairs and 27 YRI mate-pairs, after filtering on sample relatedness (see *Quality Control of HapMap and Genome of the Netherlands Data*).

#### Evaluating significance of genetic similarity in true mate-pairs

To evaluate whether genetic (dis)similarity in mate-pairs was significantly different than genetic similarity between two random individuals, we performed a permutation analysis. Specifically, we created ‘null’ (i.e., non-real) male-female pairs by randomly permuting the individuals in the true mate-pairs. Within any single permutation, we allowed for at most 1 real couple to enable faster sampling of random mate-pairs. We performed a total 1,000,000 permutations to generate a null distribution (**Figures 1** and **2**). Finally, we count the number of permutations that yield an average Qc that is the same or lower than the Qc measured in the true mate-pairs. The total number of such permutations divided by 1,000,000 is the exact p-value of the test. This permutation scheme was used to evaluate the significance of Qc as measured in common variants, all variants, and imputed HLA variants.

### Analysis of mate-pairs in the Genome of the Netherlands (GoNL) data

We repeated the same analysis in the Genome of the Netherlands data (GoNL), in the 239 mate-pairs that passed quality control. In the GoNL data, we estimated Qc in three sets of variants (**Table 1**): common biallelic variants only, all available single nucleotide variants regardless of frequency, and in all available variants (including insertions and deletions). For a fourth set of variants - imputed HLA variation - we measured genetic similarity using Pearson’s correlation (r), as the imputed variation data was phased and left no ambiguity as to how heterozygous genotypes correlated (e.g., the difference between observing the AB genotype in Sample 1 and the AB genotype in Sample 2; or observing the AB genotype in Sample 1 and the BA genotype in Sample 2). To evaluate the significance of Qc in true mate-pairs, we used the identical permutation scheme as used in the HapMap analysis and described above.

#### HLA imputation

We use SNP2HLA (http://software.broadinstitute.org/mpg/snp2hla/) [31] and a reference panel built from HLA typing performed in the Type 1 Diabetes Genetics Consortium (T1DGC) (containing 8,961 markers) [31] to impute SNPs, HLA types and amino acid substitutions across 8 classical HLA loci. For imputation, 3,256 SNPs in GoNL overlap the T1DGC reference panel data. After the MHC imputation was complete, we first performed quality control, removing samples where the total number of imputed alleles is > 2.5 (introduced by imprecision in the imputation algorithm) and removing all variants for which the imputation quality (‘info’) metric is < 0.8.

#### Correcting for population structure in the GoNL samples

As the Dutch samples are drawn from 11 of the 12 provinces in the Netherlands, subtle population structure can be observed in both common and rare variants [30]. Analysis in the original GoNL effort indicated that the first two principal components reveal a subtle north-to-south gradient, and analysis of rarer (so-called “f_2_”) variants (two alleles appearing in the entire dataset) indicate strong clustering within geographical regions (north, center, and south, as inferred by IBD analyses) [30]. We thus sought to explore whether population structure, either across the country or by province, may be confounding a potential signal for MHC-dependent mate selection. To do this, we used principal component analysis as well as province-specific analyses.

Genetic PCs are calculated on an individual basis and are an alternative means of unravelling genetic ancestral clustering between individuals. We first needed to collapse individual-level PC loadings into a single value that represented a single mate-pair. We call this collapsed PC the ‘mate-pair PC’ (PC_mp_). Assume that the PC1 loading for a female in a given mate-pair is denoted PC1_f_, and PC1 loading for the male in that mate-pair is denoted PC1m, then PC1mp (continuing up to PC ‘n’) is defined as follows:

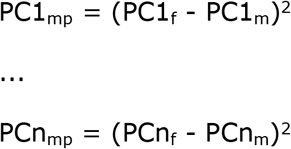

In this way, we used the PCs of the GoNL individuals to obtain, for each (real or permuted) pair of individuals, a PC_mp_ value that is equal to 0 if the loadings of the two individuals in a pair are identical for a given PC, or becomes increasingly large as the two samples’ loadings on a particular PC diverge.

We then used the mate-pairs of one random permutation of the 239 mate-pairs in GoNL to train a linear regression model that approximates the genetic similarity between two individuals (Qc), using the mate-pair PCs as defined above:

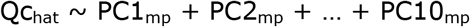

Qc_hat_ estimates the genetic similarity explained by the first 10 PCs for all mate-pairs (real or permuted) as well as residuals (Qc_res_) from this regression. If there is preferential mating among the true mate-pairs in GoNL, the residuals of this regression model should be systematically different compared to residuals from randomly-assigned male-female pairs. We performed the same initial permutation analysis, on the whole set of 239 true mate-pairs, but using Qc_res_ (instead of Qc) as a measure of genetic similarity adjusted for population stratification. We then compared where the average Qcres across the 239 true mate-pairs falls within the distribution of average Qc_res_ across 239 randomly generated male-female pairs.

### Genetic dissimilarity in non-MHC regions

In addition to permuting mate-pairs to establish a null distribution for Qc, we also wanted to establish a null distribution of Qc by randomly sampling regions from the genome that were matched to the MHC based on different characteristics. Because the MHC is an extremely unique genomic region — in gene density, in span of linkage disequilibrium, and in genetic variability — it is nearly impossible to identify regions of the genome that behave identically to the MHC. To identify genomically similar regions to the MHC from which we could construct a null distribution for Qc, we identified regions that either (1) were the same genomic span as the MHC (~3.6 Mb), or (2) contained approximately the same number of markers (~40k), regardless of the linear span of that window. For each criterion (SNP density or span), we randomly sampled 10,000 regions from the genome and computed average Qc across all 239 true mate-pairs, for each region; we compared these distributions to Qc calculated in true mate-pairs across the MHC.

## Acknowledgements

The Genome of the Netherlands Consortium (http://www.nlgenome.nl/) generated and analyzed the whole-genome sequencing data analyzed here. A complete list of the Genome of the Netherlands members and affiliations can be found here: http://www.nlgenome.nl/7page_id=28.

We thank Paul IW de Bakker for supporting MCS with funding from VIDI grant 91712354 from the Dutch Organization for Scientific Research (Nederlandse Organisatie voor Wetenschappelijk Onderzoek (NWO) - ZonMw) and for his critical review of the manuscript.

## Supporting information Legends

**SupplementaryMaterials.docx**: File containing Supplementary Figures 1 through 5, referenced in the main text.

**CoverletterPlosGenCretuStancu.docx**: Cover letter to the PLoS Genetics editors

## References

1. De Bakker PIW, McVean G, Sabeti PC, Miretti MM, Green T, Marchini J, et al. A high-resolution HLA and SNP haplotype map for disease association studies in the extended human MHC. Nat Genet. Nature Publishing Group; 2006;38: 1166–1172.

2. Horton R, Gibson R, Coggill P, Miretti M, Allcock RJ, Almeida J, et al. Variation analysis and gene annotation of eight MHC haplotypes: the MHC Haplotype Project. Immunogenetics. 2008;60: 1–18.

3. Horton R, Wilming L, Rand V, Lovering RC, Bruford EA, Khodiyar VK, et al. Gene map of the extended human MHC. Nat Rev Genet. 2004;5: 889–899.

4. Welter D, MacArthur J, Morales J, Burdett T, Hall P, Junkins H, et al. The NHGRI GWAS Catalog, a curated resource of SNP-trait associations. Nucleic Acids Res. 2014;42: D1001–6.

5. Pereyra F, Jia X, McLaren PJ, Telenti A, de Bakker PIW, Walker BD, et al. The major genetic determinants of HIV-1 control affect HLA class I peptide presentation. Science. 2010;330: 1551–1557.

6. Hinks A, Bowes J, Cobb J, Ainsworth HC, Marion MC, Comeau ME, et al. Fine-mapping the MHC locus in juvenile idiopathic arthritis (JIA) reveals genetic heterogeneity corresponding to distinct adult inflammatory arthritic diseases. Ann Rheum Dis. 2017;76: 765–772.

7. Xie G, Roshandel D, Sherva R, Monach PA, Lu EY, Kung T, et al. Association of Granulomatosis With Polyangiitis (Wegener’s) With HLA–DPB1*04 and SEMA6A Gene Variants: Evidence From Genome-Wide Analysis. Arthritis & Rheumatism. 2013;65: 2457–2468.

8. Cortes A, Pulit SL, Leo PJ, Pointon JJ, Robinson PC, Weisman MH, et al. Major histocompatibility complex associations of ankylosing spondylitis are complex and involve further epistasis with ERAP1. Nat Commun. 2015;6: 7146.

9. Zhang F-R, Liu H, Irwanto A, Fu X-A, Li Y, Yu G-Q, et al. HLA-B*13:01 and the dapsone hypersensitivity syndrome. N Engl J Med. 2013;369: 1620–1628.

10. Purcell SM, Wray NR, Stone JL, Visscher PM, O’Donovan MC, Sullivan PF, et al. Common polygenic variation contributes to risk of schizophrenia and bipolar disorder. Nature. 2009;460: 748–752.

11. Potts WK, Wakeland EK. Evolution of diversity at the major histocompatibility complex. Trends Ecol Evol. 1990;5: 181–187.

12. Penn, Penn, Potts. The Evolution of Mating Preferences and Major Histocompatibility Complex Genes. Am Nat. 1999;153: 145.

13. Potts WK, Manning CJ, Wakeland EK. Mating patterns in seminatural populations of mice influenced by MHC genotype. Nature. 1991;352: 619–621.

14. Olsson M, Madsen T, Nordby J, Wapstra E, Ujvari B, Wittsell H. Major histocompatibility complex and mate choice in sand lizards. Proc Biol Sci. 2003;270 Suppl 2: S254–6.

15. Landry C, Garant D, Duchesne P, Bernatchez L. “Good genes as heterozygosity”: the major histocompatibility complex and mate choice in Atlantic salmon (Salmo salar). Proc Biol Sci. 2001;268: 1279–1285.

16. OlsÉn KH, Grahn M, Lohm J, Langefors Å. MHC and kin discrimination in juvenile Arctic charr, Salvelinus alpinus (L.). Anim Behav. 1998;56: 319–327.

17. Chaix R, Cao C, Donnelly P. Is mate choice in humans MHC-dependent? Array, editor. PLoS Genet. Public Library of Science; 2008;4: 1–5.

18. Aeschlimann PB, Häberli MA, Reusch TBH, Boehm T, Milinski M. Female sticklebacks Gasterosteus aculeatus use self-reference to optimize MHC allele number during mate selection. Behav Ecol Sociobiol. Springer-Verlag; 2003;54: 119–126.

19. Yamazaki K, Boyse EA, Miké V, Thaler HT, Mathieson BJ, Abbott J, et al. Control of mating preferences in mice by genes in the major histocompatibility complex. J Exp Med. 1976;144: 1324–1335.

20. Penn D, Potts WK. Untrained mice discriminate MHC-determined odors. Physiol Behav. 1998;64: 235–243.

21. Brown RE, Roser B, Singh PB. Class I and class II regions of the major histocompatibility complex both contribute to individual odors in congenic inbred strains of rats. Behav Genet. 1989;19: 659–674.

22. Ober C, Weitkamp LR, Cox N, Dytch H, Kostyu D, Elias S. HLA and mate choice in humans. Am J Hum Genet. 1997;61: 497–504.

23. Hedrick PW, Black FL. HLA and mate selection: no evidence in South Amerindians. Am J Hum Genet. 1997;61: 505–511.

24. Ihara Y, Aoki K, Tokunaga K, Takahashi K, Juji T. HLA and Human Mate Choice. Tests on Japanese Couples. Anthropol Sci. 2000;108: 199–214.

25. Wedekind C, Seebeck T, Bettens F, Paepke AJ. MHC-dependent mate preferences in humans. Proc Biol Sci. 1995;260: 245–249.

26. Wedekind C, Füri S. Body odour preferences in men and women: do they aim for specific MHC combinations or simply heterozygosity? Proc Biol Sci. 1997;264: 1471–1479.

27. Frazer KA, Ballinger DG, Cox DR, Hinds DA, Stuve LL, Gibbs RA, et al. A second generation human haplotype map of over 3.1 million SNPs. Nature. 2007;449: 851–861.

28. Altshuler DM, Gibbs RA, Peltonen L, Dermitzakis E, Schaffner SF, Yu F, et al. Integrating common and rare genetic variation in diverse human populations. Nature. 2010;467: 52–58.

29. Derti A, Cenik C, Kraft P, Roth FP. Absence of Evidence for MHC–Dependent Mate Selection within HapMap Populations. PLoS Genet. Public Library of Science; 2010;6: e1000925.

30. Francioli LC, Menelaou A, Pulit SL, van Dijk F, Palamara PF, de Bakker PIW, et al. Whole-genome sequence variation, population structure and demographic history of the Dutch population. Nat Genet. 2014;46: 818–825.

31. Jia X, Han B, Onengut-Gumuscu S, Chen W-M, Concannon PJ, Rich SS, et al. Imputing amino acid polymorphisms in human leukocyte antigens. PLoS One. 2013;8: e64683.

32. The International HapMap Consortium., Gibbs RA, Belmont JW, Hardenbol P, Willis TD, Yu F, et al. The International HapMap Project. Nature. 2003;426: 789–796.

33. International HapMap Consortium. A haplotype map of the human genome. Nature. 2005;437: 1299–1320.

34. Purcell S, Neale B, Todd-Brown K, Thomas L, Ferreira M a. R, Bender D, et al. PLINK: a tool set for whole-genome association and population-based linkage analyses. Am J Hum Genet. 2007;81: 559–575.

35. Mathieson I, McVean G. Differential confounding of rare and common variants in spatially structured populations. Nat Genet. Nature Publishing Group; 2012;44: 243–246.

36. Mathieson I, McVean G. Demography and the Age of Rare Variants. PLoS Genet. 2014;10: e1004528.

37. Auton A, Abecasis GR, Altshuler DM, Durbin RM, Bentley DR, Chakravarti A, et al. A global reference for human genetic variation. Nature. 2015;526: 68–74.

38. Silventoinen K, Kaprio J, Lahelma E, Viken RJ, Rose RJ. Assortative mating by body height and BMI: Finnish twins and their spouses. Am J Hum Biol. 2003;15: 620–627.

39. Vandenburg SG. Assortative mating, or who marries whom? Behav Genet. 1972;2: 127–157.

40. Hippisley-Cox J, Coupland C, Pringle M, Crown N, Hammersley V. Married couples’ risk of same disease: cross sectional study. BMJ. 2002;325: 636.

41. Willemsen G, Vink JM, Boomsma DI. Assortative mating may explain spouses’ risk of same disease. BMJ. 2003;326: 396.

42. Sudlow C, Gallacher J, Allen N, Beral V, Burton P, Danesh J, et al. UK biobank: an open access resource for identifying the causes of a wide range of complex diseases of middle and old age. PLoS Med. 2015;12: e1001779.

43. Robinson MR, Kleinman A, Graff M, Vinkhuyzen AAE, Couper D, Miller MB, et al. Genetic evidence of assortative mating in humans. Nat hum behav. 2017;1: 0016.

44. Buhler S, Nunes JM, Sanchez-Mazas A. HLA class I molecular variation and peptide-binding properties suggest a model of joint divergent asymmetric selection. Immunogenetics. 2016;68: 401–416.

45. Chang CC, Chow CC, Tellier LC, Vattikuti S, Purcell SM, Lee JJ. Second-generation PLINK: rising to the challenge of larger and richer datasets. Gigascience. 2015;4: 1–16.

